# The autophagy activator Spermidine reduces neuroinflammation and soluble amyloid beta in an Alzheimer’s disease mouse model

**DOI:** 10.1101/2021.10.28.466219

**Authors:** Kiara Freitag, Nele Sterczyk, Benedikt Obermayer, Julia Schulz, Judith Houtman, Lara Fleck, Caroline Braeuning, Roberto Sansevrino, Christian Hoffmann, Dragomir Milovanovic, Stephan J. Sigrist, Thomas Conrad, Dieter Beule, Frank L. Heppner, Marina Jendrach

**Author notes:** contributed equally (Frank L. Heppner, Marina Jendrach).

## Abstract

Deposition of amyloid beta (Aβ) along with glia cell-mediated neuroinflammation are prominent pathogenic hallmarks of Alzheimer’s disease (AD). In recent years, impairment of autophagy has been found to be another important feature, contributing to AD progression and aging. Therefore, we assessed the effect of the autophagy activator Spermidine, a small body-endogenous polyamine often used as dietary supplement and known to promote longevity, on glia cell-mediated neuroinflammation. Spermidine reduced TLR3- and TLR4- mediated inflammatory processes in microglia and astrocytes by decreasing cytotoxicity, inflammasome activity and NF-κB signaling. In line with these anti-inflammatory effects, oral treatment of the amyloid prone AD-like APPPS1 mice with Spermidine reduced neuroinflammation and neurotoxic soluble Aβ. Mechanistically, single nuclei sequencing revealed microglia as one of the main targets of Spermidine treatment, with increased expression of genes implicated in cell motility and phagocytosis. Thus, Spermidine provides a promising therapeutic potential to target glia cells in AD progression.

## Introduction

Neuroinflammation plays an essential role in the development and progression of various neurodegenerative diseases. It belongs to the main hallmarks of Alzheimer’s disease (AD), alongside extracellular plaques containing amyloid-beta (Aβ) peptides and neurofibrillary tangles consisting of hyperphosphorylated microtubule-associated protein (MAP) tau [1]. The link between neuroinflammation and neurodegenerative diseases was further strengthened by the profound effects of maternal immune activation on the development of neurodegenerative diseases [2, 3]. Injection of the viral mimetic PolyI:C, a synthetic analog of double-stranded RNA, into wild type (WT) mice was sufficient to induce an AD-related pathology [4], further demonstrating a crucial role of inflammatory events in the initiation of a vicious cycle of neuropathological alterations. Microglia, the brain’s intrinsic myeloid cells, and astrocytes are the predominant cytokine-producing cells of the CNS. Both cell types are essential for maintaining brain homeostasis and respond to danger signals by transforming into an activated state, characterized by increased proliferation and cytokine release [1].

A growing set of data, including those derived from genome-wide association studies of various human diseases by the Wellcome Trust Case Control Consortium [5], indicates that autophagy, one of the crucial degradation and quality control pathways of the cell, can regulate inflammatory processes. Mice deficient in the autophagic protein ATG16L1 exhibited a specific increase of IL-1β and IL-18 in macrophages and severe colitis, which could be ameliorated by anti-IL-1β and IL-18 antibody administration [6]. Similar results were achieved with mice lacking the key autophagic factor MAP1LC3B after exposure to LPS or caecal ligation and puncture-induced sepsis [7]. Recently, we could show that a reduction of the key autophagic protein Beclin1 (BECN1), which is also decreased in AD patients [8, 9], resulted in an increased release of IL-1β and IL-18 by microglia [10]. The multimeric NLRP3 inflammasome complex, responsible for processing Pro-IL-1β and Pro-IL-18 into its mature forms by activated Caspase-1 (CASP1) [11], can be degraded by autophagy [10, 12]. Lack of the NLRP3-inflammasome axis induced amelioration of neuroinflammation and disease pathology in several neurodegenerative mouse models [13- 15], thus emphasizing that activation of autophagy presents an intriguing therapeutic target to counteract neuroinflammation.

The small endogenous polyamine and nutritional supplement Spermidine is known to induce autophagy by inhibiting different acetyltransferases [16, 17] and to extend the life span of flies, worms and yeast [17-20]. In addition, Spermidine supplementation improved clinical scores and neuroinflammation in mice with experimental autoimmune encephalomyelitis (EAE) [20, 21], protected dopaminergic neurons in a Parkinson’s disease rat model [22], and exhibited neuroprotective effects and anti-inflammatory properties in a murine model of accelerated aging [23]. Consistent with these observations, Spermidine decreased the inflammatory response of macrophages and the microglial cell line BV2 upon LPS stimulation *in vitro* [24-26]. Recent data showing that polyamines improved age-impaired cognitive function and tau-mediated memory impairment in mice [27, 28] and impaired COVID-19 virus particle production [29], highlighting the need to investigate the therapeutic potential of Spermidine and its molecular mechanisms also in chronic inflammation and AD pathology, which has been missing so far. Here, we can show that Spermidine interfered with key inflammatory signaling pathways *in vitro* and AD-associated neuroinflammation *in vivo*. Applying single nuclei sequencing, we identified microglial changes in the expression of genes associated with cell motility and phagocytosis as one target mechanism of Spermidine’s action in the brains of APPPS1 mice, correlating with reduced soluble Aβ levels. We therefore propose considering Spermidine as a therapeutic option for neuroinflammation and AD pathology.

## Results

### Pre-treatment with Spermidine reduced TLR4- and TLR3-mediated inflammatory response of microglia and astrocytes

We began by assessing the effects of Spermidine on the acute neuroinflammatory response of glial cells *in vitro*. First, neonatal microglia were pre-treated with Spermidine and subsequently stimulated with an established protocol for activation of the TLR4 pathway by application of LPS followed by ATP (Fig. 1a). Indeed, the presence of Spermidine reduced the LPS/ATP-induced release of IL-1β, IL-18, IL-6 and TNF-α into the cell supernatant dose-dependently as measured by ELISA, whereas IL-1β reacted most sensitively (Fig. 1b-e). Expanding this analysis by using electrochemiluminescence (MesoScale Discovery panel), we detected reduced levels of eight out of ten cytokines after Spermidine treatment (Supplementary Fig. 1a). To assess whether Spermidine exerts microglia-specific anti-inflammatory effects, the TLR4 pathway of neonatal astrocytes was activated according to the scheme depicted in Fig. 1a. Similar to microglia, astrocytes showed a dose-dependent reduction of IL-6 in the supernatant in response to Spermidine pre-incubation (Fig. 1f), while IL-1β was released at low levels (Supplementary Fig. 1b).

**Figure 1:**
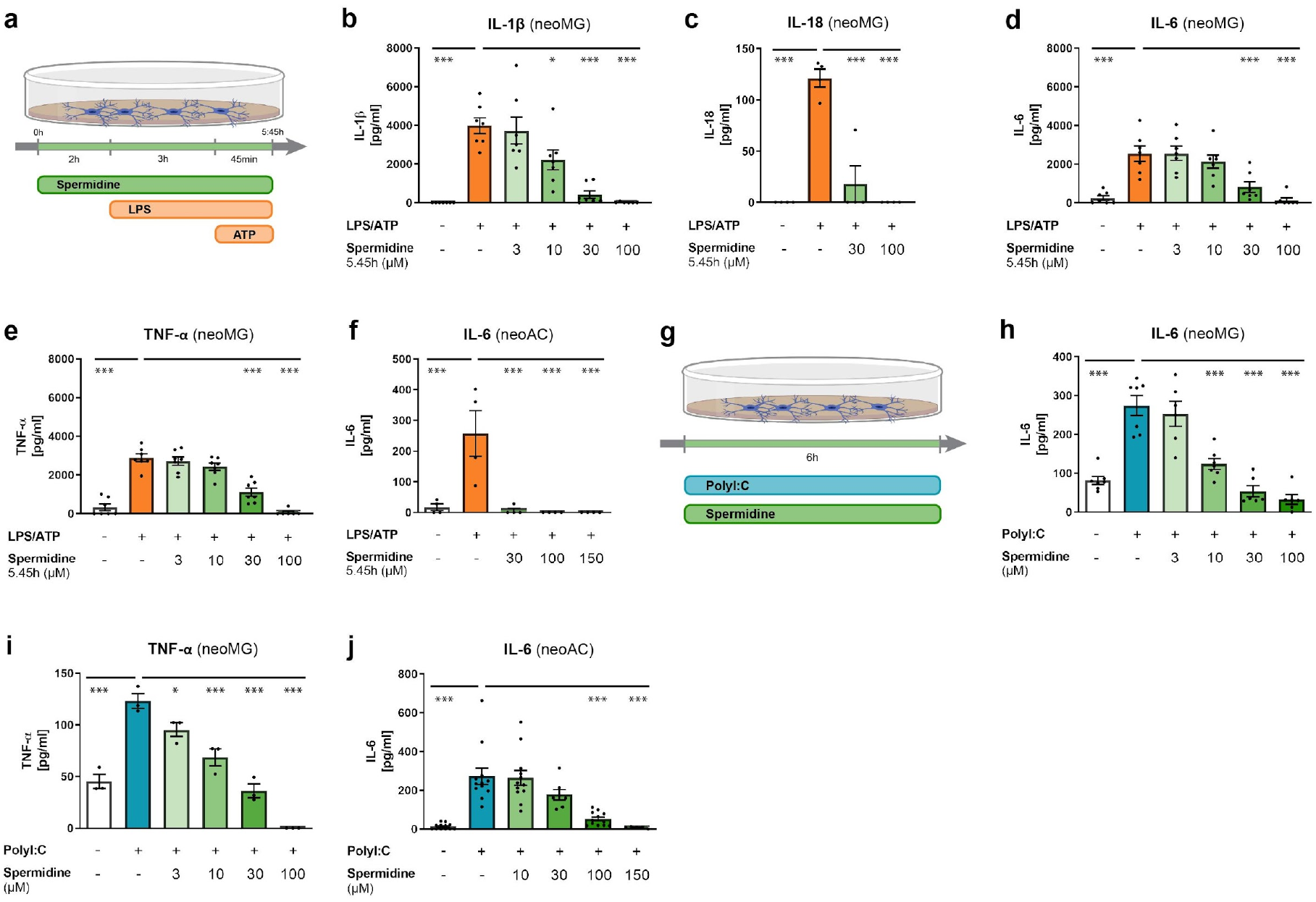
Pre-treatment with Spermidine reduced TLR4- and TLR3-mediated inflammatory response of microglia and astrocytes. (**a-f**) Neonatal microglia (neoMG) and neonatal astrocytes (neoAC) were treated with Spermidine at indicated concentrations for 5.45 h, LPS (1 µg/ml), and ATP (2 mM) as depicted in the scheme (**a**). (**b-e**) The IL-1β, IL-18, IL-6 and TNF-α concentration in the cell supernatant of neonatal microglia (neoMG) was determined by ELISA; n = 4-7. (**f**) The IL-6 concentration in the cell supernatant of neonatal astrocytes (neoAC) was determined by ELISA; n = 4. (**g-j**) Cells were treated with PolyI:C (50 µg/ml) and Spermidine at various concentrations for 6 h as depicted in the scheme (**g**). **(h)** The IL-6 concentration in the cell supernatant of neonatal microglia was determined by ELISA; n = 3-7. **(i)** The IL-6 concentration in the cell supernatant of neonatal astrocytes was determined by ELISA; n = 5-12. **(j)** The TNF-α concentration in the cell supernatant of neonatal microglia was determined by ELISA; n = 3-7. Mean ± SEM, one-way ANOVA, Dunnett’s post hoc test (reference= LPS/ATP-treated or PolyI:C-treated cells), * p < 0.05, ** p < 0.01, *** p < 0.001.

To examine whether the anti-inflammatory effects of Spermidine were limited to TLR4- mediated inflammation, we studied its effects on the TLR3 pathway by using the viral dsRNA PolyI:C (Fig. 1g). Confirming published observations [30, 31], PolyI:C treatment induced the release of IL-6 and TNF-α by neonatal microglia and IL-6 by neonatal astrocytes, as measured in the cell supernatant by ELISA (Fig. 1h-j). Similar to the observed effects on the TLR4 pathway, Spermidine treatment dose-dependently reduced the PolyI:C-induced release of IL- 6 and TNF-α in neonatal microglia and neonatal astrocytes (Fig. 1h-j), with microglia showing a higher sensitivity to Spermidine. Again, we gained a broader overview of the cytokine specificity of Spermidine by using electrochemiluminescence (MesoScale Discovery panel). Apart from IL-6 and TNF-α, the release of IL-12, IL-10, and KC/GRO was also significantly induced by PolyI:C treatment, all of which were significantly blocked by Spermidine (Supplementary Fig. 1c, d), emphasizing its broad interference spectrum beyond TLR4- mediated inflammation.

To exclude that a putative Spermidine-induced cytotoxicity interfered with the cytokine response, lactate dehydrogenase (LDH) release was measured in the cell supernatant for all stimulation schemes, showing no cytotoxic effect of the Spermidine concentrations used here, instead revealing protective effects after LPS/ATP treatment (Supplementary Fig. 1e-f).

### Pre-treatment with Spermidine reduced the glial inflammatory response by targeting the NF-κB pathway

Next, we investigated the underlying molecular mechanism behind cytokine reduction after pre-treatment with Spermidine. Transcriptional analysis by RT-qPCR revealed that Spermidine reduced the amount of LPS/ATP-induced *Il-1β, Il-6* and *Tnf-α* mRNA dose-dependently in neonatal microglia (Fig. 2a) and PolyI:C-induced gene expression of *II-6* and *Tnf-α* in both neonatal microglia and astrocytes (Fig. 2b-c). In the BV2 cell line, Spermidine has been shown to target the transcription factor NF-κB, responsible for the induction of *IL-1β, IL-6* and *TNF-α* expression (Choi and Park 2012). Therefore, we assessed NF-κB p65 phosphorylation by Western blot. In neonatal microglia, Spermidine reduced the LPS/ATP-mediated increase in total NF-κB p65 and phosphorylated NF-κB p65 significantly (Fig. 2d). In neonatal astrocytes, Spermidine reduced PolyI:C-induced phosphorylation of NF-κB p65, whilst not modulating total NF-κB levels (Fig. 2e). Taken together, Spermidine decreased TLR4- and TLR3-mediated cytokine release in neonatal microglia and neonatal astrocytes by reducing the induction of the respective mRNA expression via diminished NF-κB activity.

**Figure 2:**
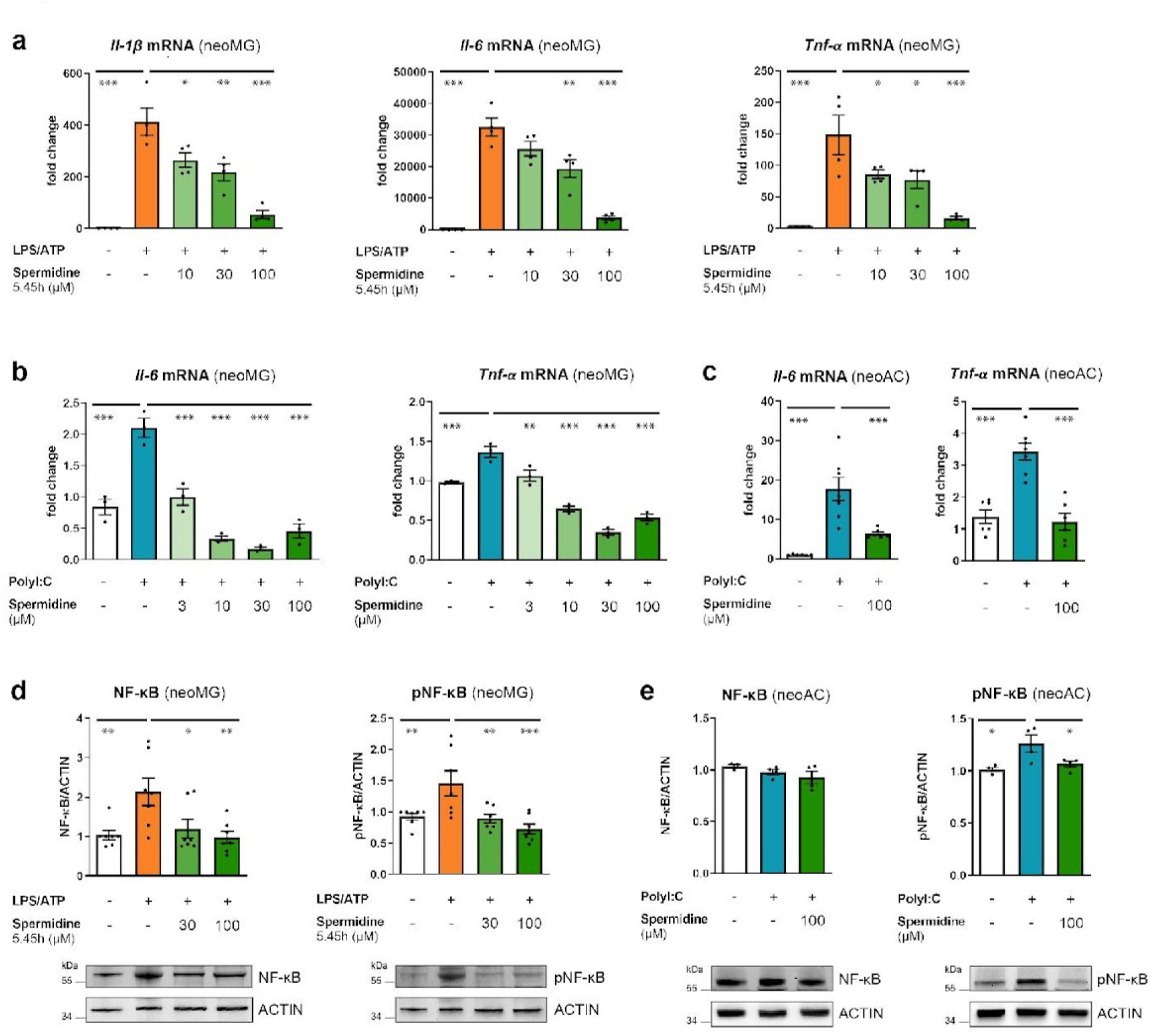
Pre-treatment with Spermidine reduced the glial inflammatory response by targeting the NF-κB pathway. Neonatal microglia (neoMG) and neonatal astrocytes (neoAC) were either treated with LPS (1 µg/ml) and ATP (2 mM) or PolyI:C (50 µg/ml) and the indicated Spermidine concentrations as depicted in Fig. 1a or Fig. 1g. **(a)** The gene expression of *Il-1β, Tnf-α* and *Il-6* was assessed by RT-qPCR after treatment of neonatal microglia as depicted in Fig. 1a. Their expression was normalized to A*ctin* and displayed as fold change compared to non-treated control cells; n = 4. **(b)** The gene expression of *Il-6* and *Tnf-α* was assessed by RT-qPCR after treatment of neonatal microglia as depicted in Fig. 1g. Its expression was normalized to *Actin* and displayed as fold change compared to non-treated control microglia; n = 3. **(c)** The gene expression of *Il-6* and *Tnf-α* was assessed by RT-qPCR after treatment of neonatal astrocytes as depicted in Fig. 1g. Its expression was normalized to *Actin* and displayed as fold change compared to non-treated control astrocytes; n = 6-7. **(d)** Levels of phosphorylated NF-κB (pNF-κB) and NF-κB were determined by Western blot in neonatal microglia treated as depicted in Fig. 1a. Representative images are shown and protein levels are displayed as fold changes compared to non-treated controls normalized to ACTIN; n = 7. **(e)** Levels of phosphorylated NF-κB (pNF-κB) and NF-κB were determined by Western blot in neonatal astrocytes treated as depicted in Fig. 1g. Representative images are shown and protein levels are displayed as fold changes compared to non-treated controls normalized to ACTIN; n = 3-4. Mean ± SEM, one-way ANOVA, Dunnett’s post hoc test (reference= LPS/ATP-treated or PolyI:C-treated cells), * p < 0.05, ** p < 0.01, *** p < 0.001.

As we have shown previously that BECN1-mediated autophagy controls IL-1β and IL-18 release by the degradation of NLRP3 and inflammasomes [10], we assessed whether pre-treatment with Spermidine also affects the inflammasome pathway. The inflammasome is a cytosolic oligomeric signaling platform, controlling IL-1β and IL-18 levels post-translationally. Their precursors, Pro-IL-1β and Pro-IL-18, are processed by activated CASP1 at the inflammasome, which is formed after LPS/ATP stimulation and consists of NLRP3 and ASC [32].

While *Nlrp3* and *Pro-Casp1* mRNA expression as well as NLRP3 protein expression were not significantly altered upon Spermidine treatment (Supplementary Fig. 1g,h), protein expression of Pro-IL-1β was reduced by administering 30 µM Spermidine (Supplementar Fig. 1i), correlating with the *Il-1β* mRNA levels (Fig. 2a). In addition, protein levels of cleaved CASP1 p20 in the supernatant were decreased, while levels of Pro-CASP1 were strongly increased after treatment with Spermidine, (Supplementary Fig. 1j, k), implying that Spermidine regulates inflammasome activity in addition to NF-κB-controlled transcription.

### Spermidine interferes with the IL-1β processing pathway in activated microglia

To further investigate the effect of Spermidine on inflammasome activity, we treated microglia with Spermidine after priming them with LPS and then added ATP afterwards (Fig. 3a). In this setting, we investigated the effects of Spermidine on previously activated microglia, thereby mimicking a therapeutic scenario.

**Figure 3:**
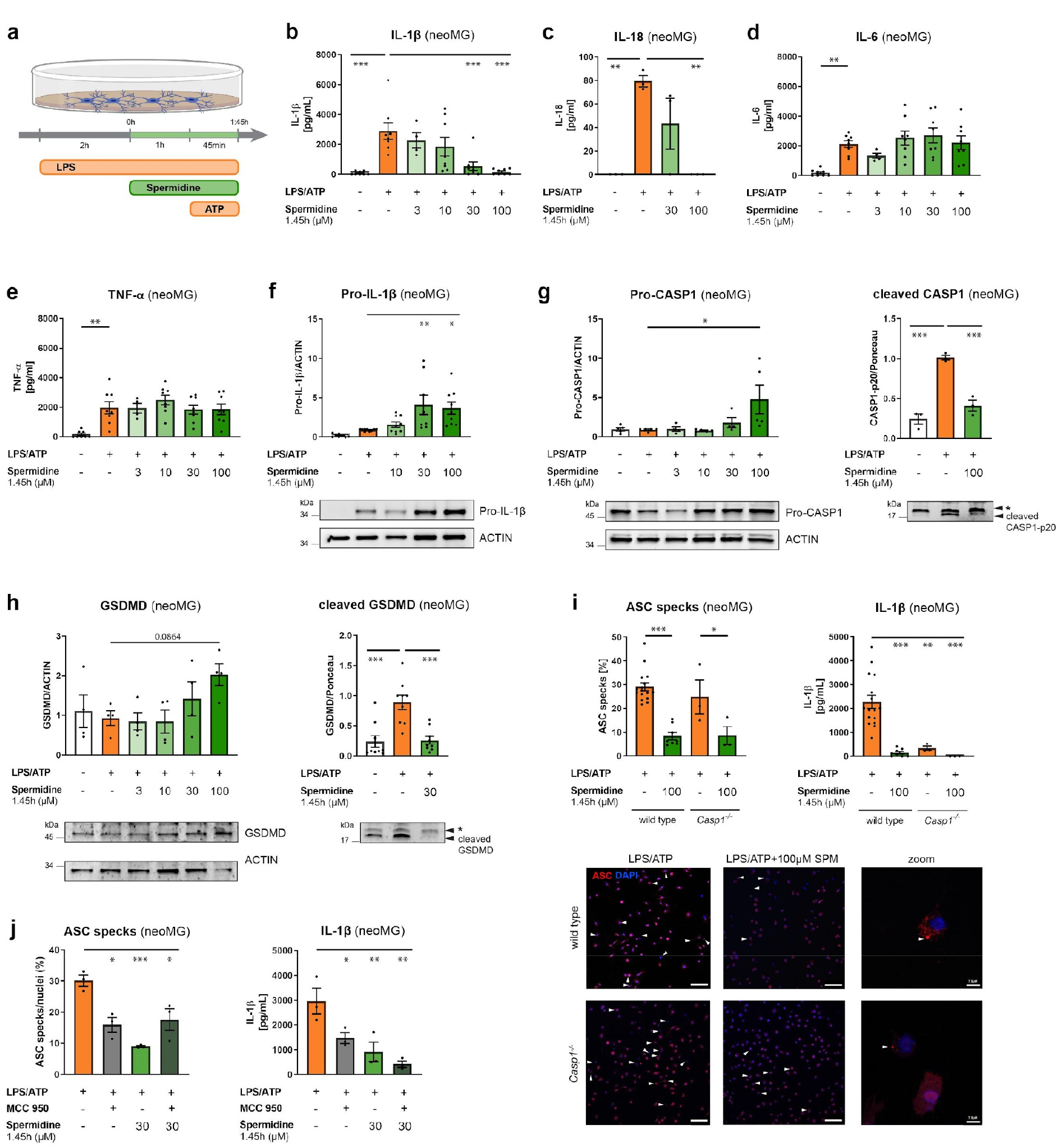
Spermidine interfered with the IL-1β processing pathway in activated microglia. Neonatal microglia (neoMG) were treated with LPS (1 µg/ml) and Spermidine at indicated concentrations for 1.45 h and ATP (2 mM) as depicted in the scheme (**a**). **(b)** The IL-1β concentration in the cell supernatant was determined by ELISA; n = 4-8. **(c)** The IL-18 concentration in the cell supernatant was determined by ELISA; n = 3. **(d)** The IL-6 concentration in the cell supernatant was determined by ELISA; n = 4-8. **(e)** The TNF-α concentration in the cell supernatant was determined by ELISA; n = 4-8. **(f)** Pro-IL-1β protein levels were determined by Western blot and normalized to ACTIN. Representative images are shown and values are displayed as fold changes compared to LPS/ATP-treated cells; n = 8-9. **(g)** Cellular Pro-CASP1 and cleaved CASP1 levels in the supernatant were determined by Western blot (* non-specific band). Pro-CASP1 was normalized to ACTIN (n = 4-8) and CASP1 was normalized on whole protein content determined by Ponceau S staining (n = 3). Values are displayed as fold changes compared to LPS/ATP-treated cells. **(h)** Cellular and cleaved GSDMD (C-terminal fragment) levels in the supernatant were determined by Western blot (* non- specific band). GSDMD was normalized to ACTIN (n = 4) and the C-terminal fragment was normalized to whole protein content determined by Ponceau S staining (n = 9). Values are displayed as fold changes compared to LPS/ATP-treated cells. **(i)** Neonatal WT and *Casp1*^-/-^ microglia were stimulated as shown in (**a**) but with 4 mM ATP to increase the number of inflammasomes. Cells were stained for ASC (red) to visualize inflammasomes and with DAPI (blue) for nuclear staining as shown in the representative images (scale bar = 100 µm). The percentage of ASC specks in respect to the number of total cells (DAPI positive cells) was determined (left). One-way ANOVA, Tukey’s post hoc test. The IL-1β concentration in the cell supernatant was assessed by ELISA (right); WT: n = 8-16; *Casp1*^-/-^: n = 3. **(j)**Neonatal microglia were stimulated as shown in (**a**) and MCC950 was added 15 min before addition of ATP. Cells were stained for ASC to visualize inflammasomes and with DAPI for nuclear staining. The percentage of ASC specks in respect to the number of total cells (DAPI positive cells) was determined (left). The IL-1β concentration in the cell supernatant was assessed by ELISA (right); n = 3. Mean ± SEM, one-way ANOVA, Dunnett’s post hoc test (reference = LPS/ATP-treated cells) if not stated otherwise, * p < 0.05, ** p < 0.01, *** p < 0.001.

Cytokine analysis showed that Spermidine reduced IL-1β and IL-18 release into the cell supernatant dose-dependently (Fig. 3b, c), while not altering the release of IL-6 and TNF-α (Fig. 3d, e). We expanded our analysis by using electrochemiluminescence assays and found that in fact only IL-1β and IL-18 out of all measured cytokines were regulated by Spermidine (Supplementary Fig. 3a). Again, Spermidine-mediated cytoprotective effects were shown by reduced LDH release in activated Spermidine-treated microglia (Supplementary Fig. 3b). The transcriptional regulation of *Il-1β, Il-6* and *Tnf-α* by Spermidine revealed no alterations (Supplementary Fig. 2c). However, increased protein levels of Pro-IL-1β were found (Fig. 3f), indicating reduced processing. Also increased levels of uncleaved Pro-CASP1 in microglia treated with 100 µM Spermidine were detected, correlating with a reduction of cleaved and activated CASP1 in the supernatant (Fig. 3g). In agreement with this observation, cleavage of Gasdermin D (GSDMD), another substrate of CASP1, was also strongly reduced in Spermidine- treated microglia (Fig. 3h).

Upon observing that the precursors of the IL-1β processing pathway accumulated while the active forms were reduced, we investigated the effects of Spermidine on the inflammasome. NLRP3 expression was not altered on the mRNA or protein level (Supplementary Fig. 2d). However, staining and quantification of ASC specks/inflammasomes revealed that Spermidine addition reduced the number of ASC specks significantly. A similar reduction was also detected in *Casp1*^-/-^ microglia (Fig. 3i), indicating that Spermidine did not directly interfere with Pro- CASP1 cleavage but rather with inflammasome formation. To test this hypothesis, the ASC- oligomerization inhibitor MCC950 [33] was added before addition of ATP. As no additive effects of MCC950 to the Spermidine-mediated effects with regard to IL-1β release or number of ASC specks could be detected, we deduce that Spermidine is indeed interfering with ASC- oligomerization and with the inflammasome formation in activated microglia (Fig. 3j). Consistent with this hypothesis, Western blot analyses for ASC after chemical crosslinking showed reduced appearance of ASC oligomers in Spermidine-treated cells (Supplementary Fig. 2e), while the amount of ASC monomers was not altered (Supplementary Fig. 2f). Thus, Spermidine treatment of activated microglia reduced IL-1β processing by interfering with the oligomerization of ASC-positive inflammasomes, elucidating a novel regulatory mechanism of Spermidine in addition to targeting NF-κB-mediated transcription of pro-inflammatory genes.

Taken together, while Spermidine-mediated protective effects were shown in neurons, dendritic cells, macrophages and BV2 cells [25, 34-37], our results indicate that Spermidine has the potential to target various glial populations in the brain as well.

### Autophagy activation by Spermidine modulated the inflammatory response of glial cells

Several beneficial effects of Spermidine are attributed to Spermidine-mediated induction of autophagy [16, 17]. Thus, we assessed whether the Spermidine-mediated anti-inflammatory effects in glial cells can be linked to autophagy. Spermidine treatment significantly upregulated the key autophagic proteins LC3-II and BECN1 in LPS/ATP- and PolyI:C-treated microglia and PolyI:C-treated astrocytes (Supplementary Fig. 3a, b). In agreement with a previous study, LC3-I was only weakly expressed by primary microglia [15]. Since the key transcription factor TFEB, that controls autophagosomal and lysosomal biogenesis [38, 39], has been found to be regulated by Spermidine in B cells [40], *Tfeb* mRNA expression was assessed. Indeed, while LPS/ATP and PolyI:C significantly reduced *Tfeb* mRNA in neonatal microglia and astrocytes, Spermidine preserved *Tfeb* levels comparable to non-treated cells (Supplementary Fig. 3c).

Next, we determined whether autophagy induction is crucial for the anti-inflammatory action of Spermidine. Here, either full medium or amino acid-free starvation medium HBSS, known to achieve a robust induction of autophagy, was used. As shown previously [10], cell starvation impaired the release of cytokines upon LPS/ATP or PolyI:C treatment significantly, underlining that autophagy induction reduced the inflammatory response of glial cells. In addition, no further reduction of cytokine release after Spermidine treatment became discernable (Supplementary Fig. 3d,e).

Previous data showed that impairment of autophagy reversed several beneficial effects of Spermidine [16-18, 41]. Therefore, cells were treated according to the scheme in Fig. 3a and the autophagy-inhibitor 3-MA was added simultaneously with Spermidine. Addition of 3-MA abolished the Spermidine-mediated reduction of IL-1β release completely (Supplementary Fig. 3f). Furthermore, we genetically excised the *Becn1* gene by treating neonatal microglia derived from BECN1^flox/flox^.CX3CR1^CreERT2^ mice with Tamoxifen or ethanol as a control, ultimately resulting in a strong reduction of BECN1 protein expression in Tamoxifen-treated cells (Supplementary Fig. 3g). LPS/ATP-treated BECN1-deficient microglia showed a significantly decreased sensitivity to Spermidine with regard to IL-1β, IL-6 and TNF-α release (Supplementary Fig. 3h). Taken together, these data confirm that Spermidine induces autophagy in glial cells, thus mediating the reduction of cytokine release.

### Spermidine treatment reduced progressive neuroinflammation in APPPS1 mice

In order to further assess Spermidine’s anti-inflammatory potential, microglia were isolated from adult WT mice and treated with the established stimulation schemes shown in Fig. 1 and Fig. 3. Similarly to neonatal microglia, post-LPS treatment with Spermidine reduced the release of IL-1β (Supplementary Fig. 4a) and pre-treatment with Spermidine blocked the LPS/ATP-induced release of IL-1β, IL-6 and TNF-α (Supplementary Fig. 4b) and PolyI:C-induced release of IL-6 and TNF-α (Supplementary Fig. 4c). Expanding our analysis to an *in vivo*-like chronic inflammatory setting, APPPS1 mice harboring transgenes for the human amyloid precursor protein (APP) bearing the Swedish mutation as well as presenilin 1 (PSEN1) were used, which develop a strong Aβ pathology including neuroinflammation [42]. Acute whole hemisphere slice cultures derived from WT mice or 200 day old APPPS1 mice were pre-treated with Spermidine and subsequently stimulated with LPS and ATP (Fig. 4a). LPS/ATP treatment of APPPS1 slice cultures revealed a massive release of IL-1β and IL-6 compared to slices from WT mice. Spermidine treatment reduced the IL-1β and IL-6 release of slice cultures of both genotypes (Fig. 4a), suggesting that Spermidine might influence neuroinflammation in APPPS1 mice *in vivo*.

**Figure 4:**
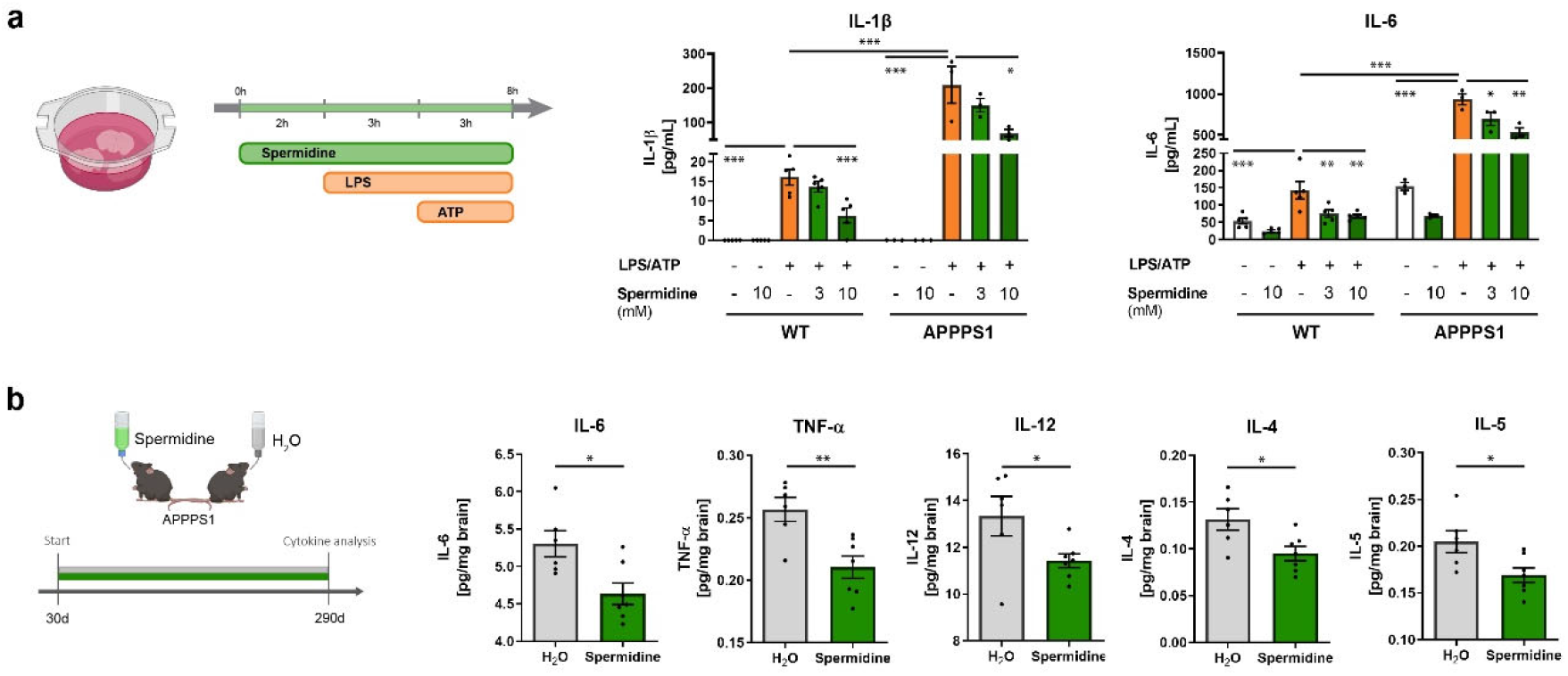
Spermidine treatment reduced progressive neuroinflammation in APPPS1 mice. **(a)** Hemispheres of WT and APPPS1 mice were coronally sliced and treated with the indicated Spermidine concentration, LPS (10 µg/ml) and ATP (5 mM) as depicted in the scheme. The IL-1β and IL-6 concentration in the supernatant was determined by ELISA. One-way ANOVA, Dunnett’s post hoc test (reference = LPS/ATP-treated cells); n = 3-5, two-way ANOVA. **(b)** APPPS1 mice were treated with 3 mM Spermidine via their drinking water starting at 30 days (d) until mice reached an age of 290 days according to the depicted treatment scheme. Spermidine-treated APPPS1 mice were compared to non- treated controls (H_2_O). The content of the cytokines IL-6, TNF-α, IL-12, IL-4 and IL-5 in the TBS fraction of brain homogenates of 290d old male Spermidine-treated mice and water controls was measured using electrochemiluminescence (MesoScale Discovery panel). APPPS1 H_2_O (n = 6), APPPS1 Spermidine (n = 7). Mean ± SEM, two-tailed t-test, * p < 0.05, ** p < 0.01, *** p < 0.001.

Thus, APPPS1 mice were treated with 3 mM Spermidine [27] via their drinking water starting prior to disease onset (namely substantial Aβ deposition), at the age of 30 days (Fig. 4b). Compared to control APPPS1 mice that received water (H_2_O), Spermidine-supplemented animals showed no differences in fluid uptake per day (Supplementary Fig. 4d). To examine the neuroinflammatory status of 120 day and 290 day old male Spermidine-treated APPPS1 mice, ten cytokines were quantified by electrochemiluminescence in brain homogenates containing soluble proteins. Spermidine supplementation significantly reduced the AD-relevant pro-inflammatory cytokines IL-6, TNF-α, IL-12, IL-4 and IL-5 in 290 day old mice (Fig. 4b), whilst not altering IL-1β (combined measurement of Pro- IL-1β and IL-1β), IFN-γ, IL-2, IL- 10 and KC/GRO or neuroinflammation at 120 days (Supplementary Fig. 4e, f), confirming the anti-inflammatory potential of Spermidine.

### Spermidine treatment of APPPS1 mice induced transcriptomic alterations in microglia

To gain insights into the underlying mechanisms behind these changes and the cell populations affected by Spermidine treatment *in vivo*, we performed comparative single nuclei sequencing (snRNA-seq) using hemispheres of male Spermidine-treated APPPS1 mice, H_2_O APPPS1 controls as well as WT mice in triplicates at 180 days, representing a midpoint in pathology (Fig. 5a). Using fluorescence-activated cell sorted single nuclei and a 10x Genomics platform (Supplementary Fig. 5a), we detected between 6500 and 10000 cells per mouse at a median depth of 1400-1700 genes. Automated clustering revealed 34 clusters which we grouped into 7 major cell types, including neurons, oligodendrocytes, microglia, oligodendrocyte precursors (OPC), astrocytes, macrophages and fibroblasts/ vascular cells, using label transfer from a previously published mouse brain reference dataset (https://doi.org/10.1101/2021.04.25.441313) (Fig. 5b, c).

**Figure 5:**
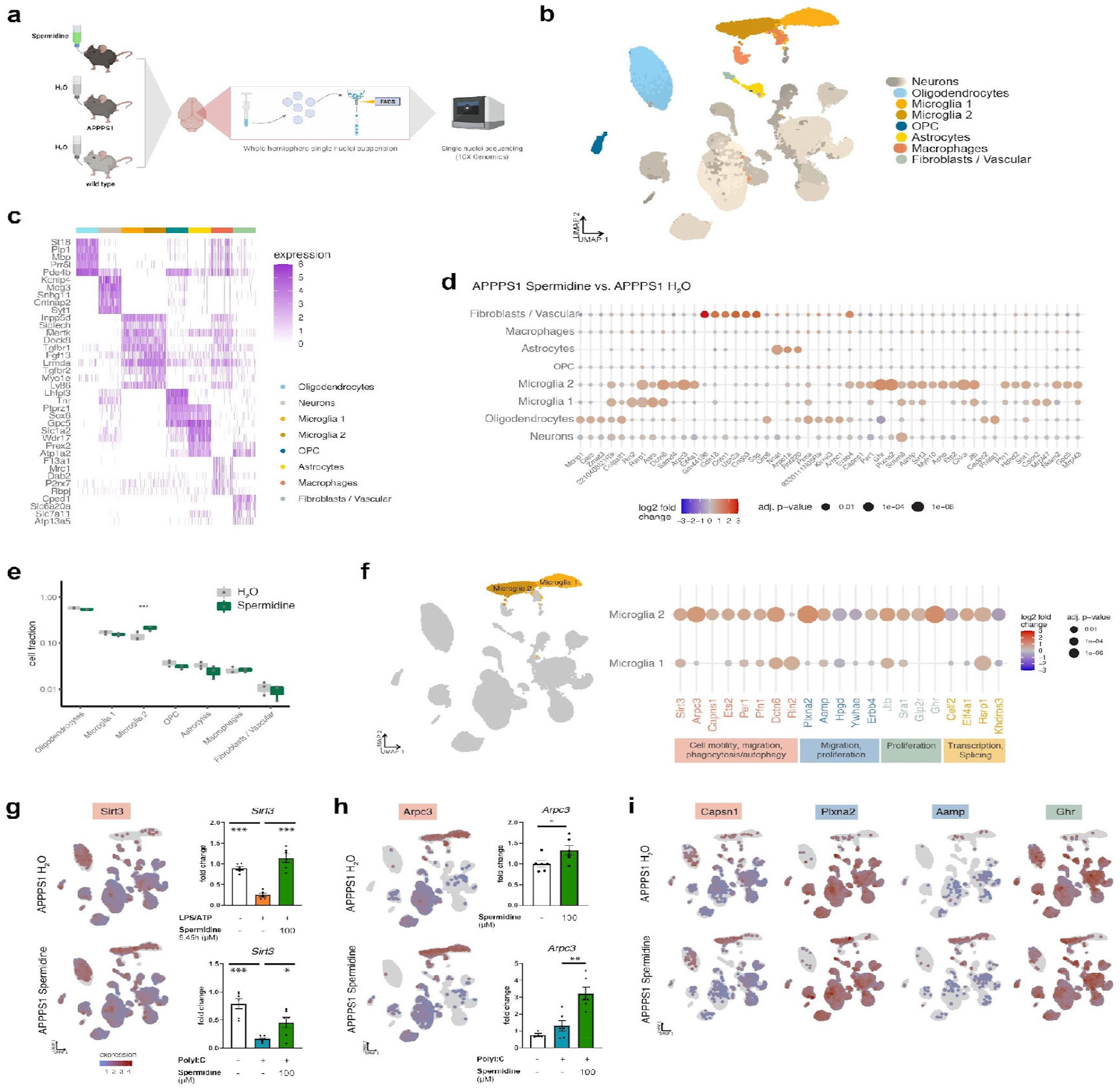
Spermidine treatment of APPPS1 mice induced transcriptomic alterations in microglia. **(a)** Single nuclei sequencing of hemispheres harvested from Spermidine-treated APPPS1, H_2_O APPPS1 and H_2_O control mice was performed of FACS-sorted DAPI-stained nuclei using the 10x Genomics platform (n = 3). **(b)** UMAP embedding and clustering of the snRNAseq data, together with annotation of major cell types. **(c)** Heatmap showing the top 5 marker genes for 300 cells in each of the major cell types. **(d)** Dot plot for the top 50 genes in a cell-type-specific differential expression analysis between Spermidine-treated APPPS1 and H_2_O APPPS1 mice. Color scale indicates log2 fold change, dot size indicates adjusted p-value. **(e)** Cluster abundance in Spermidine-treated APPPS1 and H_2_O APPPS1 mice. P-values from mixed-effects binomial model. * p < 0.05, ** p < 0.01, *** p < 0.001 **(f)** Same as d, for selected genes differential in microglia 1 or 2. Associated pathways are color-coded. (**g-h**) left panels: Expression of *Sirt3* and *Arpc3* in Spermidine-treated APPPS1 (top) and H_2_O APPPS1 mice (bottom). Color scale indicates normalized expression, grey dots represent no data. right panels: Neonatal microglia were treated with the indicated concentrations of Spermidine in combination with LPS (1 µg/ml) and ATP (2 mM) or with PolyI:C (50µg/ml) and the gene expression was assessed by RT-qPCR. Their expression was normalized to A*ctin* and displayed as fold change compared to non-treated control cells; n = 4-6, one-way ANOVA, Dunnett’s post hoc test. Mean ± SEM, * p < 0.05, ** p < 0.01, *** p < 0.01. **(i)** As in (g-h) for additional genes.

In agreement with previous single cell transcriptomic analyses of APPPS1 mice [43, 44], we detected two microglia subpopulations. The microglia 2 cluster appeared only in APPPS1 but not in WT mice, thus presenting an AD-associated activated microglia phenotype largely equivalent to the classical disease-associated microglia published by Keren-Shaul et al. [43] (Supplementary Fig. 5b-e). To characterize the main characteristics of both microglia clusters, differential gene expression followed by gene set enrichment analysis between these populations was performed. Compared to cluster 1, the AD-associated cluster 2 revealed a downregulation of genes involved in phagocytosis, endocytosis, cell adhesion and cell polarity, while upregulating neuroinflammatory responses, cell-cycle transition and autophagy (Supplementary Fig. 5f).

By focusing on the changes induced by Spermidine treatment of APPPS1 mice, we found the strongest transcriptional changes in microglia. Fewer genes were altered in oligodendrocytes, neurons, and astrocytes, while OPC and macrophages remained almost unaffected (Fig. 5d, Supplementary Fig. 5g, h). Interestingly, we observed that Spermidine significantly increased the abundance of microglia cluster 2 (Fig. 5e). Thus, genes differentially expressed in Spermidine-treated compared to H_2_O APPPS1 mice were specifically assessed in microglia clusters 1 and 2 (Fig. 5f). We found the following anti-inflammatory-associated genes to be regulated: *Pfn1* [45], *Glp2r* [46], *Per1* [47] and *Sirt3*. The upregulation of *Sirt3* by Spermidine in the AD-associated microglia cluster 2 was especially striking (Fig. 5g), as the NAD-dependent deacetlyase SIRT3 is known to exhibit anti-inflammatory effects, targeting several cytokines including the NLRP3 inflammasome in the IL-1β processing pathway [48, 49]. By RT-qPCR analysis, we could confirm that activation of neonatal microglia induced a reduction in *Sirt3* levels, which was rescued by Spermidine treatment, thus making SIRT3 a potential driver of the observed Spermidine-mediated effects (Fig. 5g).

Notably, among the top differentially expressed genes in microglia were genes associated with cell motility, cell migration (*Arpc3, Capns1, Pfn1, Plxna2, Aamp, Erbb4, Ywhae, Hpgd*), phagocytosis (*Arpc3, Capns1, Pfn1, Dctn6, Rin2*), autophagy (*Arpc3, Capns1, Ets2, Per1, Ghr*), proliferation (*Aamp, Ets2, Erbb4, Hpgd, Glp2r, Ghr, Jtb, Sra1, Ywhae*), transcription and alternative splicing (*Celf2, Eif4a1, Rsrp1, Khdrbs3*) (Fig. 5f-i). The increased abundance of microglia cluster 2 could be connected to the induction of proliferation-associated genes by Spermidine. Accordingly, gene set enrichment analysis revealed the following Gene Ontology terms to be significantly regulated by Spermidine: glial cell migration, microtubule organization center localization, cell matrix adhesion and the semaphorin plexin signaling pathway. Indeed, Spermidine treatment of neonatal microglia increased expression of the actin nucleation gene *Arpc3* (Fig. 5h) and reduced the levels of the anti-proliferatory gene *Hpgd* (Supplementary Fig. 5i), matching the snRNA-seq findings. Notably, Spermidine might exert some of its function not solely by affecting the transcriptome but also the proteome, as indicated by our *in vitro* findings.

### Spermidine treatment altered phagocytic behavior of microglia and reduced soluble Aβ in APPPS1 mice

Based on current literature, microglia in early AD pathology are characterized by acute activation including enhanced proliferative, phagocytic and migratory behavior, which ceases and reverts during the progression of the disease and the chronic activation of microglia [50, 51]. The characterization of the AD-associated cluster 2 indicates that these APPPS1 microglia seem to be in the progressed state during which endocytosis and phagocytosis are downregulated (Supplementary Fig. 5f). However, Spermidine appears to rescue the impairment of phagocytosis and endocytosis. We therefore hypothesize that Spermidine prolongs the early activated state of microglia characterized by increased phagocytosis, cell motility, migration and proliferation, thus maintaining the surveillance mode of microglia and potentially reducing Aβ. In line with this hypothesis, Spermidine treatment reduced the levels of the transcriptional regulator Celf2 (Fig. 5f), which negatively regulates the phagocytic receptor TREM2 [52] and increased the expression of the phagocytic receptors *Trem2* and *Cd36* in activated neonatal microglia *in vitro* (Fig. 6a, b).

**Figure 6:**
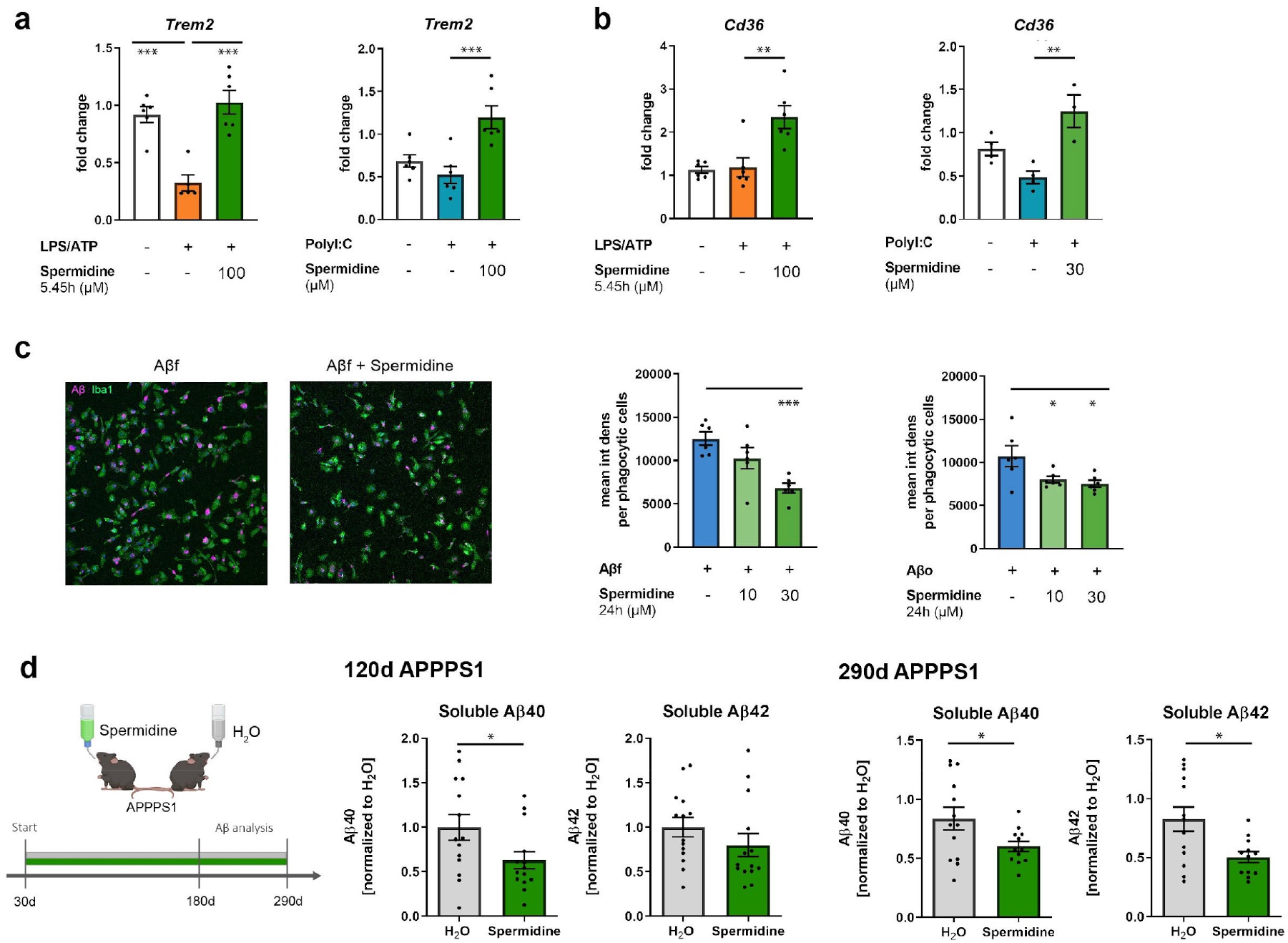
Spermidine treatment altered phagocytic behavior of microglia and reduced soluble Aβ in APPPS1 mice. (**a-b**) Neonatal microglia were treated with 100 µM Spermidine in combination with LPS (1 µg/ml) and ATP (2 mM) or with PolyI:C (50µg/ml) and the gene expression of *Trem2* and *Cd36* was assessed by RT-qPCR. Their expression was normalized to A*ctin* and displayed as fold change compared to non-treated control cells; n = 5-6. **(c)** Neonatal microglia were pre-treated with the indicated concentrations of Spermidine and fluorescently labelled (pink) fibrillary (Aβf) or oligomeric (Aβo) was added for 24 h. Microglia cells were stained with Iba1 (green). The mean intensity density per phagocytic cells was assessed by confocal microscopy. Representative images are shown; n = 6, one-way ANOVA, Dunnett’s post hoc test (reference = Aβ-treated cells). **(d)** APPPS1 mice were treated with 3 mM Spermidine via their drinking water starting at 30 days (d) until mice reached an age of 120 days or 290 days according to the depicted treatment scheme. Spermidine-treated APPPS1 mice were compared to non-treated controls (H_2_O). The Aβ40 and Aβ42 content was measured in the TBS (soluble) fraction of brain homogenates of (**b**) 120d old or (**c**) 290d old Spermidine-treated mice and water controls using electrochemiluminescence (MesoScale Discovery panel). Values were normalized to water controls. 120d APPPS1 H_2_O (n = 14), 120d APPPS1 Spermidine (n = 14), 290d APPPS1 H_2_O (n = 14), 290d APPPS1 Spermidine (n = 12). Mean ± SEM, two-tailed t-test, * p < 0.05, ** p < 0.01, *** p < 0.01.

To test whether Spermidine directly modifies the phagocytic behavior, neonatal microglia were pre-treated with Spermidine before addition of oligomeric and fibrillary fluorescently labelled Aβ for 24 h. Spermidine treatment significantly decreased the mean Aβ signal per phagocytic cell (Fig. 6c), while the percentage of phagocytic cells was not altered (Supplementary Fig. 6a), indicating that Spermidine enhances the degradation of Aβ. A histogram, showing the intensity signal of each phagocytic cell, revealed that the amount of cells with low Aβ signal was increased upon Spermidine treatment, while a lower percentage with high fluorescent signal was found, supporting the idea of increased Aβ degradation in Spermidine-treated cells (Supplementary Fig. 6b).

Finally, Aβ pathology was analyzed in Spermidine-treated APPPS1 mice over time at 120 and 290 days, when pathology is known to have reached a plateau [42] (Fig. 6d). After consecutive protein extractions, soluble and insoluble Aβ40 and Aβ42 levels were measured by electrochemiluminescence (MesoScale Discovery panel). In line with the snRNA-seq findings and the *in vitro* data, Spermidine supplementation significantly reduced soluble Aβ40 levels in both 120 day old and 290 day old APPPS1 mice by 40 % and soluble Aβ42 in 290 day old mice by 49 % (Fig. 6d), whilst not affecting insoluble Aβ (Supplementary Fig. 6c). These findings were further substantiated as no differences in insoluble Aβ plaque covered area or plaque size after staining tissue sections with the fluorescent dye pFTAA were observed (Supplementary Fig. 6d). As APP processing was not affected by Spermidine treatment (Supplementary Fig. 6e), we conclude that Spermidine treatment targeted soluble Aβ by maintaining microglial phagocytic and migratory behavior.

## Discussion

Preventing AD or at least delaying disease progression still presents an urgent unmet clinical need. Based on recent advances in our understanding of AD pathogenesis that resulted in the appreciation of the impact of neuroinflammation and autophagy, we assessed the therapeutic effects of the autophagic activator Spermidine on the main cytokine-producing glial cells, namely microglia and astrocytes. By modeling neuroinflammation *in vitro* through activation of the TLR3 and TLR4 pathway, we found that Spermidine treatment abolished the release of pro-inflammatory cytokines by microglia and astrocytes by targeting various pathways: NF- κB-mediated transcription of cytokines, cytotoxicity and the assembly of the NLRP3 inflammasome. In agreement with previous studies [16, 17], Spermidine-induced autophagy in glial cells accounts for the observed reduction of neuroinflammation. In line with our *in vitro* findings, we observed a substantial reduction of inflammation in brain slice cultures as well as in 290 day old male APPPS1 mice treated with Spermidine *in vivo*. Consistent with these findings, pro-inflammatory cytokine production was decreased and disease pathology was ameliorated in Spermidine-treated EAE mice, a murine model for multiple sclerosis [20, 21].

Interestingly, single nuclei sequencing of a hemisphere from 180 day old Spermidine-treated APPPS1 mice revealed microglia as the main cytokine-producing cell type targeted by Spermidine. On the contrary, astrocytes were barely affected, despite the *in vitro* data showing anti-inflammatory responses after Spermidine treatment in both cell types, which could be explained by their decreased sensitivity to Spermidine shown *in vitro*. Among the significantly regulated candidate genes in microglia was the NAD-dependent deacetylase *Sirt3*. Similarly to Spermidine, SIRT3 was shown to exhibit anti-inflammatory effects, specifically targeting the NLRP3 inflammasome and IL-1β processing pathway as well as IL-6 and TNF-α [48, 49], indicating that Spermidine might exert some of its effects by modulating SIRT3. In addition, SIRT3 was shown to be decreased in AD patients and in microglial BV2 cells upon Aβ treatment [53, 54] and to promote the migratory behavior of microglia [55], thus targeting all pathways shown to be affected by Spermidine treatment. Of note, clear anti-inflammatory effects of Spermidine became only apparent at 290 days, when APPPS1 mice show neuroinflammation changes. Thus, only a few differentially expressed genes related to inflammation could be found at 180 days, correlating with the lack of anti-inflammatory effects of Spermidine at 120 days.

While changes in *Sirt3* and other genes with anti-inflammatory properties (*Pfn1, Glp2r, Per1)* that were up-regulated in 180 day old Spermidine-treated APPPS1 mice might pave the path for the reduction in neuroinflammation observed at 290 days, our *in vitro* analyses on the NLRP3 inflammasome pathway as well as recent publications on a post-translational modification called hypusination [27, 28], indicate that Spermidine might exert some of its function also on the post-translational level. Among those might be autophagic proteins, which are mainly regulated post-translationally.

The most profound effects of Spermidine on the transcriptome were seen in the AD-associated microglia cluster 2, which was characterized by increased migration, cell motility, phagocytosis, autophagy and cell proliferation. While acute activation of microglia in early disease pathology induces microglial phagocytosis and migration towards plaques, later stages of AD pathology and chronic priming of microglia with Aβ have adverse effects [50, 51]. In accordance, microglial motility in the presence of Aβ plaques was found to be decreased in APPPS1 mice compared to control mice when a focal lesion was performed by a laser [56]. By promoting genes known to be involved in cell motility, migration, phagocytosis and autophagy, Spermidine seems to delay the late stage AD-associated microglial phenotype. In addition, Spermidine also increased the abundance of microglia cluster 2. Although it is still under discussion whether proliferation of microglia in AD is beneficial or detrimental [57], we conclude that Spermidine mediates an enlargement of a microglial population characterized by increased phagocytosis and cell motility. Several regulated genes as *Arpc3* [58], *Glp2r* [59], *Sirt3* [53] and *Per1* [60, 61] have been shown to exert protective effects in neurodegenerative diseases or reverse memory deficits in various models, underlining the observed protective effects of Spermidine.

In line with the snRNA-seq findings, Spermidine treatment also significantly reduced Aβ *in vitro* as well as soluble Aβ levels at 120 days and at 290 days, while Aβ plaque burden and size were not altered. These results correlate with recent data by De Risi et al. [62], who observed that Spermidine decreased soluble Aβ and α-synuclein in a mouse model with mild cognitive impairment. Currently, it is thought that soluble Aβ causes more synaptotoxicity than plaque-bound insoluble Aβ by altering synaptic transmission and mediating synaptic loss and neuronal death, thus stressing the importance of targeting soluble Aβ in AD [63-65]. In 120 d old mice only Aβ40, was reduced, while at the later time point both Aβ species were less present in the soluble fraction. Interestingly, studies have shown that soluble Aβ40 has a substantial pathogenetic effect on neuronal survival underlining its qualitative impact in AD [66-68]. In addition, very recent data imply that microglia create core plaques as a protective measure in order to shield the brain from soluble Aβ [69], thus presenting an additional mechanism of Spermidine-mediated mitigation of AD pathologic progression. As the APPPS1 mouse model exhibits a fast disease progression with a strong genetically-driven amyloid pathology appearing at 60 days [42], the substantial effect of Spermidine on soluble Aβ supports the potential of Spermidine to counteract or at least slow AD progression. Additionally, neuroinflammation is a known driver for plaque formation [2, 3], thus the anti-inflammatory effects of Spermidine might potentially affect plaque size in APPPS1 mice older than the analyzed 290 day old animals.

In agreement with our findings, several studies suggest an involvement of autophagic mechanisms in regulating AD. Similarly to Spermidine, strong activators of autophagy such as fasting or caloric restriction, Rapamycin, an inhibitor of the mechanistic Target of Rapamycin (mTOR), and Metformin were found to prolong the life span of several species and to reduce Aβ deposition in different mouse models [70-73], emphasizing that the herein described effects of Spermidine in APPPS1 mice on Aβ burden are genuine. Spermidine’s multifarious intracellular interference points are advantageous, particularly in light of its good tolerability and its uncomplicated oral administration method. Therefore, we consider the body- endogenous substance Spermidine as a novel, attractive therapeutic dietary supplement in AD as it attenuated AD-relevant neuroinflammation and reduced synaptotoxic soluble Aβ. Since Spermidine supplementation is already tested in humans, the extension of Spermidine supplementation from individuals with subjective cognitive decline [28, 74-76] to clinical trials aimed at testing Spermidine efficacy in AD patients appears justified.

## Materials and Methods

### Mice and Spermidine treatment

APPPS1^+/−^ mice [42] were used as an Alzheimer’s disease-like mouse model. *Casp1*^-/-^ mice were a kind gift from F. Knauf and M. Reichel, Medizinische Klinik m.S. Nephrologie und Internistische Intensivmedizin, Charité Berlin. Beclin1^flox/flox^ mice were a kind gift from Tony Wyss-Coray (Stanford University School of Medicine/USA).

APPPS1^+/−^ mice and littermate wild type control mice were treated with 3 mM Spermidine dissolved in their drinking water (changed twice a week) from an age of 30 days until an age of either 120 days or 290 days. Control mice received only water (H_2_O). Prior to each exchange of the drinking bottles, the weight of the bottles was determined and used to calculate the average volume consumed per animal per day. Animals were kept in individually ventilated cages with a 12 h light cycle with food and water *ad libitum*. All animal experiments were conducted in accordance with animal welfare acts and were approved by the regional office for health and social service in Berlin (LaGeSo).

### Tissue preparation

Mice were anesthetized with isoflurane, euthanized by CO_2_ exposure and transcardially perfused with PBS. Brains were removed from the skull and sagitally divided. The left hemisphere was fixed with 4 % paraformaldehyde for 24 h at 4°C and subsequently immersed in 30% sucrose until sectioning for immunohistochemistry was performed. The right hemisphere was snap-frozen in liquid nitrogen and stored at -80°C for a 3-step protein extraction using buffers with increasing stringency as described previously [77]. In brief, the hemisphere was homogenized in Tris-buffered saline (TBS) buffer (20 mM Tris, 137 mM NaCl, pH = 7.6) to extract soluble proteins, in Triton-X buffer (TBS buffer containing 1% Triton X-100) for membrane-bound proteins and in SDS buffer (2% SDS in ddH_2_O) for insoluble proteins. The protein fractions were extracted by ultracentrifugation at 100,000 g for 45 min after initial homogenization with a tissue homogenizer and a 1 ml syringe with G26 cannulas. The respective supernatants were collected and frozen at -80°C for downstream analysis. Protein concentration was determined using the Quantipro BCA Protein Assay Kit (Pierce) according to the manufacturer’s protocol with a Tecan Infinite® 200 Pro (Tecan Life Sciences).

### Quantification of Aβ levels and pro-inflammatory cytokines

Aβ40 and Aβ42 levels of brain protein fractions were measured using the 96-well MultiSpot V-PLEX Aβ Peptide Panel 1 (6E10) Kit (MesoScale Discovery, K15200E-1). While the TBS and TX fraction were not diluted, the SDS fraction was diluted 1:500 with Diluent 35. Cytokine concentrations were measured in the undiluted TBS fraction using the V-PLEX Pro-inflammatory Panel 1 (MesoScale Discovery, K15048D1). For all samples, duplicates were measured and concentrations normalized to BCA values.

### Histology

Paraformaldehyde-fixed and sucrose-treated hemispheres were frozen and cryosectioned coronally at 40 µm using a cryostat (Thermo Scientific HM 560) and stored afterwards in cryoprotectant (0.65 g NaH2PO4 × H2O, 2.8 g Na2HPO4 in 250 ml ddH2O, pH 7.4 with 150 ml ethylene glycol, 125 ml glycerine) at 4°C until staining. For immunohistochemistry, sections were washed with PBS and incubated with pentameric formyl thiophene acetic acid (pFTAA, 1:500, Sigma-Aldrich) for 30 min at RT. Subsequently, cell nuclei were counterstained with DAPI (1:2000, Roche, 10236276001) and sections embedded in fluorescent mounting medium (Dako, S3023). For quantification of pFTAA positive Aβ plaques, images of 10 serial coronal sections per animal were taken with an Olympus BX53 microscope, equipped with the QImaging camera COLOR 12 BIT and a stage controller MAC 6000 (1.25x objective). Images were analyzed using ImageJ by defining the cortex as the region of interest. Images were converted to grey scale and by using the same threshold for all sections, the pFTAA-positive area [in %] was obtained. Additionally, the average plaque size was determined and further analyzed by performing a plaque size distribution using thresholds for the size of the pFTAA-positive particles. The average of all 10 sections per animal was displayed in the graphs.

### Brain slice culture

The brains of C57Bl/6J and APPPS1 mice were harvested, the cerebellum removed and the hemispheres mounted on a cutting disk using a thin layer of superglue. Hemispheres were cut using the Vibratome platform submerged in chilled medium consisting of DMEM medium (Invitrogen, 41966-029) supplemented with 1% penicillin/streptomycin (Sigma, P0781-20ML). Coronal slicing was performed from anterior to posterior after discarding the first 1 mm of tissue generating 10 × 300 µm sequential slices per brain with vibrating frequency set to 10 and speed to 3. Brain slices were cultured in pairs in 1 ml culture medium at 35 °C, 5 % CO_2_ in 6-well plates. Pre-treatment with the indicated Spermidine concentrations was started immediately for 2 h. Subsequently, LPS (10 µg/ml) was added to the medium for 3 h followed by the addition of ATP (5 mM) for an additional 3 h. Afterwards, the culture medium was frozen for subsequent analyses.

### Isolation and culture of adult microglia

Adult microglia were isolated from the hemispheres of 160 day old C57BL/6J mice by magnetic activated cell sorting (MACS). The manufacturer’s protocols were followed. In brief, mice were transcardially perfused with PBS and tissue dissociated with the Neural Tissue Dissociation kit (P) (Miltenyi Biotec, 130-092-628) in C-tubes (Miltenyi, 130-096-334) on a gentleMACS Octo Dissociator with Heaters (Miltenyi Biotec, 130-096-427). Afterwards, the cell suspension was labelled with CD11b microbeads (Miltenyi Biotec, 130-093-634) and passed through LS columns (Miltenyi Biotec, 130-042-401) placed on a OctoMACS™ manual separator. Subsequently, microglia were collected by column flushing and cultured in DMEM medium (Invitrogen, 41966-029) supplemented with 10% FCS (PAN-Biotech, P40-37500) and 1% penicillin/streptomycin (Sigma, P0781-20ML). Medium was changed every three days until adult microglia were treated as indicated after 8 days *in vitro* (DIV).

### Cell culture of neonatal microglia and astrocytes

Newborn mice (1-4 days old) were sacrificed by decapitation. Mixed glial cultures were prepared as described previously [10]. In brief, brains were dissected, meninges removed and brains mechanically and enzymatically homogenized with 0.005% trypsin/EDTA. Cells were cultured in complete medium consisting of DMEM medium (Invitrogen, 41966-029) supplemented with 10% FCS (PAN-Biotech, P40-37500) and 1% penicillin/streptomycin (Sigma, P0781-20ML) at 37°C with 5% CO_2_. From 7 DIV on, microglia proliferation was induced by adding 5 ng/ml GM-CSF (Miltenyi Biotec, 130-095-746) to the complete medium. Microglia were harvested at 10-13 DIV by manually shaking flasks for 6 min. Cells were treated after a settling time of 24 h. Neonatal BECN1^flox/flox^.CX3CR1^CreERT2^ microglia were treated with (Z)-4- Hydroxytamoxifen (Sigma #7904) after 5 DIV and assessed 7 days after Tamoxifen treatment. After isolating neonatal microglia, neonatal astrocytes were separated by MACS. Neonatal astrocytes were detached with 0.05% trypsin, pelleted by centrifugation and incubated with CD11b microbeads (Miltenyi Biotec, 130-093-634) for 15 min at 4°C to negatively isolate astrocytes. Afterwards, the cell suspension was passed through LS columns (Miltenyi Biotec, 130-042-401) placed on an OctoMACS™ manual separator and the flow-through containing the astrocytes was collected. Subsequently, astrocytes were cultured in complete medium for 2 days before being treated. For all experiments, 100,000 cells were seeded on 24 well plates if not stated otherwise.

### Cell treatment

For pro-inflammatory stimulation, cells were either treated with LPS (1 μg/ml, Sigma, L4391- 1MG) for 3 h followed by ATP (2 mM, Sigma Aldrich, A6419-5g; 4 mM for ASC speck/inflammasome formation) for 45 min or with PolyI:C (50 µg/ml, InVivoGen, tlrl-picw- 250) for 6 h if not stated otherwise. Spermidine trihydrochloride (Sigma, S2501-5G) diluted in complete medium was added as indicated. Autophagy was activated by keeping cells in HBSS for 2 h prior to treatment (24020-091, Invitrogen) and blocked by addition of 3-MA (Sigma-Aldrich, M9282, final concentration 10 mM). The ASC oligomerization inhibitor MCC950 (inh-mcc, Invivogen) was used with a final concentration of 300 nM.

### Western blot

For ASC crosslinking, 1 mM DSS (Thermo, A39267) was added to freshly harvested microglia in PBS for 30 min. All cell pellets were lysed in protein sample buffer containing 0.12 M Tris-HCl (pH 6.8), 4% SDS, 20% glycerol, 5% β-mercaptoethanol. Proteins were separated by Tris-Tricine polyacrylamide gel electrophoresis (PAGE) according to [78] and transferred by wet blotting onto a nitrocellulose membrane. After blocking with 1% skim milk in Tris-buffered saline with 0.5% Tween20 (TBST), the following primary antibodies were used: BECN1 (1:500, Cell Signaling, 3495), LC3 (1:500, Sigma, L8918), CASP1 and pro-CASP1 (1:500, Abcam, ab179515), IL-1β and pro-IL-1β (1:500, eBioscience, 88701388), NLRP3 (1:500, AdipoGen, AG- 20B-0014), Gasdermin D (1:500, Adipogene, AG-25B-0036-C100), ASC (1:500, Adipogene, AG- 25B-0006-C100) and ACTIN (1:10,000, Sigma, A1978). For signal detection the SuperSignal West Femto Maximum Sensitivity Substrate (ThermoFisher, 34096) was used. Western blots were analyzed by quantifying the respective intensities of each band using ImageJ. All samples were normalized to ACTIN levels or whole protein content in the supernatant.

### Quantitative real-time PCR

For total RNA isolation, the RNeasy Mini kit (Qiagen, 74104) was used and cells were directly lysed in the provided RLT lysis buffer in the cell culture plate. Reverse transcription into cDNA was performed using the High-Capacity cDNA Reverse Transcription kit (ThermoFisher, 4368813). The manufacturer’s instructions for both kits were followed. Quantitative PCR was conducted on a QuantStudio 6 Flex Real-Time PCR System (Applied Biosystems) using 12 ng cDNA per reaction. Gene expression was analyzed in 384 well plates by using the TaqMan Fast Universal Master Mix (Applied Biosystems, 4364103) and TaqMan primers for *β-Actin* (ThermoFisher, Mm00607939_s1), *Arpc3* (ThermoFisher, Mm07799871_m1), *Casp1* (ThermoFisher, Mm00438023_m1), *Cd36* (ThermoFisher, Mm01135198_m1) *Hpgd* (ThermoFisher, Mm00515121_m1), *Il-1β* (ThermoFisher, Mm00434228_m1), *Il-6* (ThermoFisher, Mm00446190_m1), *Nlrp3* (ThermoFisher, Mm00840904_m1), *Sirt3* (ThermoFisher, Mm00452131_m1), *Tfeb* (ThermoFisher, Mm00448968_m1), *Tnf-α* (ThermoFisher, Mm00443258_m1), *Trem2* (ThermoFisher, Mm04209424_g1). Within the Double delta Ct method, values were normalized to the house keeping gene *Actin* and non-treated controls.

### ELISA

Cytokine concentrations in the supernatant of cultured cells were measured using the IL-1β (eBioscience, 88701388), IL-18 (Thermo Fisher, 88-50618-22), TNF-α (eBioscience, 88723477) and IL-6 (eBioscience, 88706488) enzyme-linked immunosorbent assay (ELISA) kit according to manufacturer’s instructions. The absorption was read at a wavelength of 450 nm and a reference length of 570 nm with the microplate reader Infinite® 200 Pro (Tecan Life Sciences) and analyzed using the Magellan Tecan Software.

### Cytotoxicity assay

Cell cytotoxicity was detected by measuring lactate dehydrogenase (LDH) levels using the CytoTox 96® Non-Radioactive Cytotoxicity Assay (Promega, G1780). The manufacturer’s instructions were followed and the absorbance reflecting the LDH content in the cell supernatant was measured at 492 nm (600 nm reference) with an Infinite® 200 Pro (Tecan Life Sciences) plate reader. All values were normalized to non-treated controls.

### Immunocytochemistry and confocal microscopy

Cells were seeded at a density of 50,000 cells per well on 12 mm coverslips. After treatment, cells were fixed with 4% paraformaldehyde for 20 min, permeabilized with 0.1% Triton X-100 in PBS for another 20 min and blocked with 3% bovine serum in PBS for 1 h. The primary antibodies (anti-ASC, AdipoGen, AG-25B-0006, 1:500; anti-Iba1, Wako 019-19741, 1:1000) was added overnight at 4°C. Subsequently, cells were incubated with the fluorescent secondary antibodies (Alexa Fluor 568-conjugated anti-rabbit IgG, 1:500, Invitrogen, A11011; 488-conjugated anti-rabbit IgG, Invitrogen A21206) for 3 h at room temperature. Cell nuclei were counterstained with DAPI (Roche, 10236276001) and coverslips embedded in fluorescent mounting medium (Dako, S3023). Images were acquired using Leica TCS SP5 confocal laser scanning microscope controlled by LAS AF scan software (Leica Microsystems, Wetzlar, Germany). Z-stacks were taken and images presented as the maximum projection of the z-stack. The number and size of ASC specks was assessed using ImageJ software as described before [10].

### A® preparation

#### Labeling

A® 1-42 peptides (Cayman Chemicals) were resuspended in hexafluoroisopropanol to obtain 1 mM solution, evaporated and stored as aliquots. For each preparation, 125 µg of amyloid-® was dissolved in 2 µL DMSO, sonicated for 10 min in the waterbath and supplemented with 3x molar excess of NHS-ester ATTO647N dye (Sigma) in 1x PBS (phosphate buffer saline, Gibco) and pH was adjusted to 9 with sodium bicarbonate. After 1 h of labeling reaction in the dark at room temperature, the labeled peptides were separated using spin columns (Mobicol, Mobitec) and loaded with 0.7 mL of Sephadex G25 beads (Cytiva). Clean, chromatography-grade H_2_O (LiChrosolv LC-MS grade, Merck) was used for washing, equilibration and elution. Peptide concentrations were determined using 15% SDS-PAGE gels and comparing the band intensities of the input with the eluted fractions.

#### Maturation

Labelled A® peptides were matured according to Stine et al. (PMID: 20967580). In short, to obtain oligomeric forms, A® was resuspended in the final concentration of 1x PBS and incubated at 4°C overnight. For obtaining A® fibrils, the labelled peptides were resuspended in the final concentration of 1 mM HCl and incubated at 37°C overnight.

### Phagocytosis assay

Neonatal microglia seeded on coverslips were pre-treated for 2 h with Spermidine. 0.5 µM fibrillary and oligomeric 647-labelled Aβ was added and after 24 h cells were fixed and counterstained with Iba-1. Quantification of Z-stacks taken at the confocal microscope was performed with Image J: a mask was created for each Iba-1 stained cell body and the intensity of the Aβ signal in every cell was determined. The mean intensity/phagocytic cell was calculated as well as the number of Aβ-containing/phagocytic cells.

### Nuclei preparation

Mouse hemispheres were harvested from male mice at the age of 180 days and immediately snap frozen in liquid nitrogen and stored at -80°C until further processing for nuclei isolation. Nuclei were isolated from a single mouse hemisphere in 2 ml of pre-chilled EZ PREP lysis buffer (NUC-101, Sigma) using a glass Dounce tissue grinder (D8938, Sigma) (25 strokes with pastel A and 25 strokes with pastel B) followed by incubation for 5 minutes on ice with additional 6 ml of EZ PREP buffer. During incubation, 1 µM DAPI was added to the homogenate and subsequently filtered through a 35 µm strainer. Intact nuclei were sorted with a BD FACSAriaIII with a 70 µm configuration into 1.5 ml-Eppendorf tubes with 40 µl of 4 % BSA in PBS and RiboLock RNase Inhibitor (25 U/µl, EO0381, ThermoFisher). A FSC/SSC based gate was used to exclude debris followed by exclusion of damaged nuclei in a DAPI-A/DAPI-H (see Supplementary Fig. 5a). 150,000 events were sorted for each sample. The concentration of sorted nuclei was determined based on brightfield images and DAPI fluorescence using a Neubauer counting chamber and a Leica DMi8 microscope.

### Single nuclei sequencing

Single nuclei libraries were generated according to the Chromium Next GEM Single Cell 3′ Reagent Kits v3.1 User Guide (CG000204) by 10x Genomics. Briefly, a droplet emulsion was generated in a microfluidic chip followed by barcoded cDNA generation inside these droplets. Purified and amplified cDNA was then subjected to library preparation and sequenced on a NovaSeq 6000 instrument (Illumina) to a median depth of 45-90k reads per cell.

### Single nuclei sequencing analysis

Sequencing libraries were processed with CellRanger (v3.1.0) against the mouse genome (mm10) augmented by intronic sequences and the APP and PS1 transgenes, followed by background removal with CellBender (v0.2.0; REF doi: 10.1101/791699). snRNA-seq data analysis was performed in R (v3.6.3) with Seurat (v3.2.1) [79]. Cells with at least 250 but less than 6000 genes and less than 10% mitochondrial RNA content were combined from each library, and clustering and UMAP embedding was computed based on log normalized gene counts, with total RNA content regressed out during scaling and using 20 PCA components. Cluster annotation was aided by marker expression and label transfer with Seurat’s TransferData workflow using a previously published mouse brain dataset as reference (https://doi.org/10.1101/2021.04.25.441313). We compared cluster abundances between conditions using a mixed-effects binomial model and the lme4 package (v1.1-23). Differential gene expression was performed using DESeq2 (v1.26.0; REF PMID 25516281) on aggregated “pseudo-bulk” counts for each cluster from each sample. Pathway analysis was performed using tmod [80] (v0.44) and Gene Ontology terms from the msigdbr package (v7.2.1).

### Data analysis

All values are presented as mean ± SEM (standard error of the mean). For pairwise comparison between two experimental groups, the student’s t-test was used. Statistical differences between more than two groups were assesses with One-way ANOVA using the indicated *post hoc* test. Statistically significant values were determined using the GraphPad Prism software and are indicated as follows: *P < 0.05, **P < 0.01 and ***P < 0.001

## Acknowledgement

This work was supported by the Deutsche Forschungsgemeinschaft (DFG, German Research Foundation) under Germany’s Excellence Strategy—EXC-2049—390688087, as well as under JE-278/6-1 to MJ and SFB TRR 43, SFB TRR 167 and HE 3130/6-1 to F.L.H., by the German Center for Neurodegenerative Diseases (DZNE) Berlin, and by the European Union (PHAGO, 115976; Innovative Medicines Initiative-2; FP7-PEOPLE-2012-ITN: NeuroKine). We are grateful to Alexander Haake and Julia Bertram for excellent technical support. Treatment images were created with Biorender.com. Computation has been performed on the HPC for Research cluster of the Berlin Institute of Health.

## Author contributions

KF, SS, FH and MJ designed experiments; KF, NS and JH treated and analyzed the mice; KF, NS and LF performed experiments and analyzed data for the TLR3 pathway and astrocytes; JS and MJ performed experiments and analyzed data for the TLR4 pathway and the phagocytosis; KF, NS, JS and MJ assessed autophagy; RS, CH, and DM prepared the chemically-defined A® for *in vitro* phagocytosis assays. KF and BO prepared the figures. BO performed sequencing data analysis. All authors wrote, revised and approved the manuscript.

